# Multiple regions of primate orofacial sensorimotor cortex encode bite force and gape

**DOI:** 10.1101/2020.08.16.252817

**Authors:** Fritzie I. Arce-McShane, Barry J. Sessle, Yasheshvini Ram, Carrie A. Balcer, Callum F. Ross, Nicholas G. Hatsopoulos

## Abstract

The precise control of bite force and gape is vital for effective breakdown and manipulation of food inside the oral cavity during feeding. Yet the role of the orofacial sensorimotor cortex (OSMcx) in the control of bite force and gape is still largely unknown. The aim of this study was to elucidate how individual neurons and populations of neurons in multiple regions of OSMcx differentially encode bite force and gape when subjects *(Macaca mulatta)* generated different levels of bite force at varying gapes. We examined neuronal activity recorded simultaneously from three microelectrode arrays implanted chronically in the primary motor (MIo), primary somatosensory (SIo), and cortical masticatory (CMA) areas of OSMcx. We used generalized linear models to evaluate encoding properties of individual neurons and utilized dimensionality reduction techniques to decompose population activity into components related to specific task parameters. Individual neurons encoded bite force more strongly than gape in all three OSMCx areas although bite force was a better predictor of spiking activity in MIo versus SIo. Population activity differentiated between levels of bite force and gape while preserving task-independent temporal modulation across the behavioral trial. While activation patterns of neuronal populations were comparable across OSMCx areas, the total variance explained by task parameters was context-dependent and differed across areas. These findings suggest that the cortical control of gape may rely on computations at the population level whereas the strong encoding of bite force at the individual neuron level allows for the precise and rapid control of bite force.

**Significance Statement:** Biting a piece off an apple requires precise sensorimotor control and coordination of bite force and gape by multiple brain regions. The cortical representations of bite force and gape by individual neurons and large populations of neurons across connected motor and somatosensory areas in orofacial cortex is unknown. Here we showed that bite force was more strongly encoded than gape by individual neurons in primary motor, somatosensory, and cortical masticatory areas. Moreover, bite force was more effectively represented in motor versus somatosensory cortices. At the population level, bite force and gape were distinguishable particularly when gape was randomized from trial-to-trial. The results are important for understanding neurophysiological processes underlying masticatory dysfunctions that may occur in aging, stroke, and Alzheimer’s disease.

## Introduction

Primate feeding relies on the coordination of tongue and jaw movements and the precise control of the generation of tongue and bite forces at varying distances of jaw depression, i.e., gape, during chewing and swallowing. Bite force control is important for intra-oral breakdown of food into a bolus that is safe to swallow and easy to digest while minimizing the probability of tooth breakage and excessive tooth wear. Likewise, gape has to be controlled to accommodate ingestion and manipulation of food by the lips, tongue, and teeth during ingestion, chewing, bolus transport and swallowing. Indeed, the wide range of disorders and dysfunctions affecting the feeding system pose significant challenges for human health and enjoyment of life, including tooth loss, masticatory dysfunctions, dysphagia, neuralgia, and pain states such as temporomandibular disorders (1–7). A part of the cerebral cortex termed the orofacial sensorimotor cortex (OSMcx) is crucial for controlling orofacial sensorimotor functions, yet despite the importance of feeding behavior for human health and well-being, little is known about the role of the OSMcx in the control of bite force and gape. This limited knowledge hampers our ability to leverage the full potential of OSMcx for the development of therapies and treatments and also constrains our understanding of the role of OSMcx in feeding system evolution.

The OSMcx, which includes the primary motor (MIo), primary somatosensory (SIo), and cortical masticatory (CMA) areas, plays a crucial role in the control of complex oral sensorimotor behaviors so as to effect functionally critical, coordinated movements such as those associated with feeding and speech (7–9). Several decades of research using intracortical microstimulation (ICMS), receptive field (RF) mapping, multi-electrode array recordings, and ablative procedures suggest that these three areas play important roles in these behaviors. For example, ICMS in MIo, SIo, or CMA can evoke relatively simple movements of orofacial muscles (e.g., jaw opening, tongue protrusion) as well as more complex movements such as chewing and swallowing (10–15). Neurons in MIo and SIo have been shown to modulate their activity during feeding and performance of orofacial tasks such as the generation of tongue-protrusive force or bite force, to encode the direction and magnitude of tongue-protrusive force, to form coherent networks within and across these areas in a reciprocal manner, and to undergo learning-induced plasticity (16–22). Many of these neurons have orofacial mechanosensitive RFs and the sensory inputs from their RFs are used to modulate bite and tongue forces (10, 11, 23–26). In addition, a role for OSMcx in orofacial motor control is indicated by studies showing that reversible cold-block or ablation of OSMcx disrupts various elements of feeding performance (27–33).

While these findings in animals indicate an important role for OSMcx in the control of biting and related orofacial motor behaviors, it is unknown how functionally diverse neuronal populations in three different cortical areas (i.e., MIo, SIo, and CMA) might encode gape and bite force because activity from these areas has not been recorded simultaneously when both bite force and gape parameters are controlled. Here we present new data on the role of OSMcx of macaque monkeys in the control of two critically important behavioral variables in the mammalian orofacial feeding system: bite force and gape. The aim of this study was to elucidate how individual neurons and population of neurons in multiple regions of OSMcx differentially encode bite force and gape when subjects *(Macaca mulatta)* generated different levels of bite force at varying gapes.

## Results

Two naïve monkeys were trained to perform a behavioral task that approximates a natural feeding behavior of generating different levels of bite force at varying gapes (Fig. 1). The bite force plate was computer-controlled to open at one of three gapes prior to the start of a behavioral trial. The bite plates remained in that configuration for the entire length of the trial and returned to their initial closed configuration by the end of the trial. Strain gauges glued to the bite plates recorded the bite force. For each trial, the required bite force level was cued by the position of a target shown on a computer screen placed in front of the monkey (Fig. 1b). We used nine combinations of required bite force (3 levels) and gape (3 distances) as trial types. With training, monkeys successfully generated the required bite force levels for each of the three gapes (Fig. 1c). We recorded the bite force generated by the monkeys while simultaneously recording neuronal responses from the OSMcx areas (Fig. S1). Spiking activity of single neurons in MIo, SIo, and CMA was dynamically modulated during task performance; neurons exhibited increases and decreases in firing rates relative to the onset of bite force. Activity of some task-modulated neurons exhibited more robust tuning to gape than to bite force (compare Fig. 2a-c, top vs. bottom row) while others showed activity that varied more with bite force than with gapes (Fig. 2d-f, bottom vs. top row).

**Figure 1.**
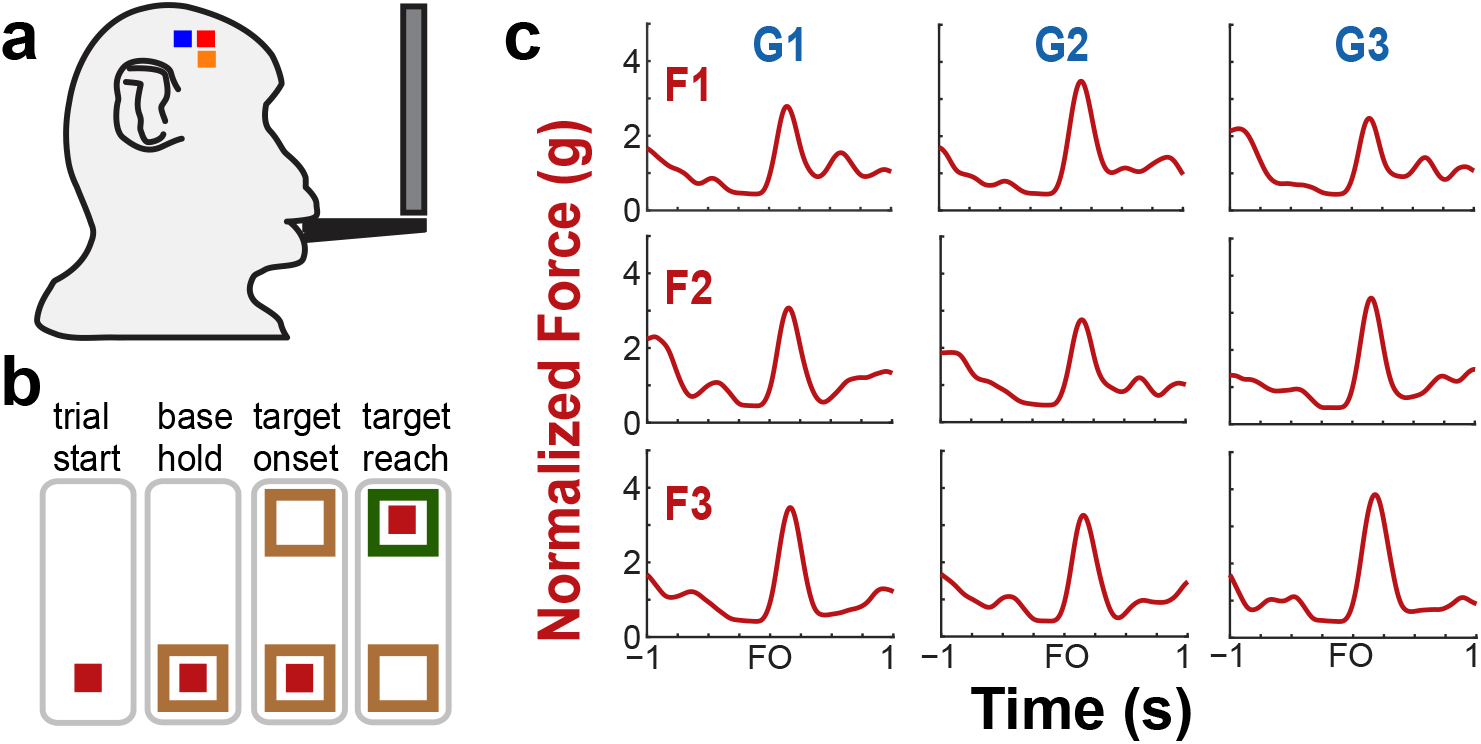
Behavioral task and performance. (**a**), Diagram of the bite force task apparatus. (**b**), Sequence of events in a trial of the bite force task. The light green square represents the force cursor while the brown and green boxes represent the base and force targets. The animals were presented with one of three target positions at one of three gapes in each trial. (**c**), Example bite force profiles of each trial type based on gape and required bite force level. Shown for monkeys H as mean bite force, across all trials of a trial type, during ±1 s relative to force onset (FO). G1, G2, G3 correspond to increasing gape distances whereas F1, F2, F3 correspond to increasing levels of required bite force.

**Figure 2.**
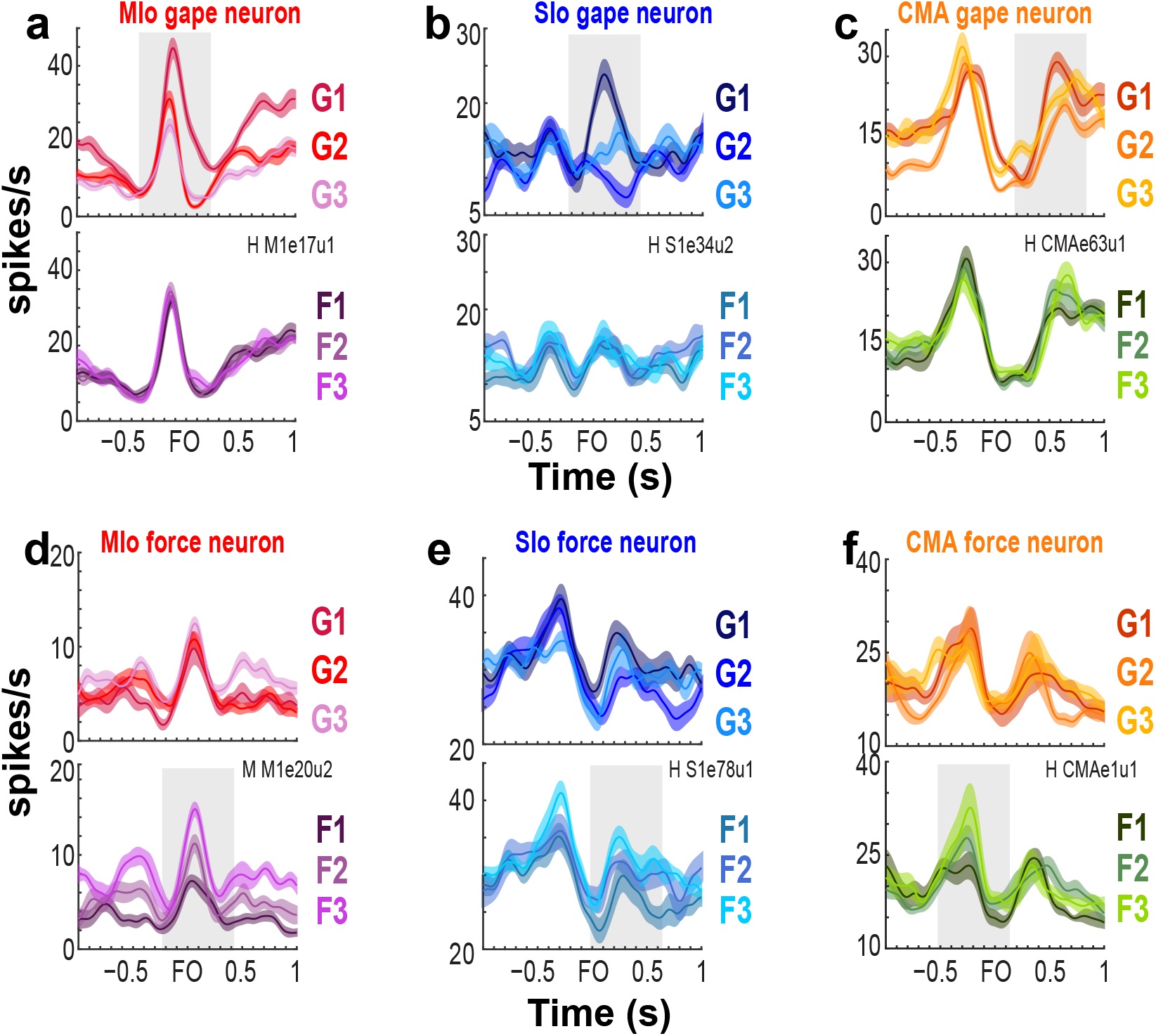
Modulation of spiking activity of single units in MIo, SIo, and CMA during task performance. (**a-c**), Peri-event time histograms (PETHs and ± 1 SE, smoothed by a 50-ms Gaussian kernel) of individual gape-related neurons simultaneously recorded in MIo, SIo, and CMA, respectively. To illustrate whether a neuron is gape-related or force-related, trials used to plot the PETHs were grouped according to gape (top row) or required bite force levels (bottom row). For example in (a), MIo gape neuron shows modulation of peak activity with different gapes (top row) but not with different levels of force (bottom row). (**d-f**), As in a-c, but for neurons whose spiking activity varied more with the varying degrees of bite force than gape.

### Encoding model

To determine the relative importance of bite force and gape in predicting the firing of neurons and to compare encoding properties among these three cortical areas, we used generalized linear models (GLM) to predict the time-varying spiking activity of each neuron. The GLM approach allows us to measure how well a model predicts the mean spike count in a small time-window based on a set of input features that included extrinsic covariates (i.e., bite force, gape, and their interaction) as well as intrinsic, spike history covariates (Fig. S2, see Methods). Predictive power was assessed by computing the area under the receiver operating characteristic curve (AUROC) on cross-validated test data (see Methods). Figure 3a-b provides an example illustrating the actual firing rates of a MIo neuron, its predicted rates based on a full encoding model that included all input features, and the AUROC for a specific cross-validation of test trials for this neuron. The encoding model predicted the spiking activity of this neuron with a mean AUROC value of 0.84 across all ten runs of cross-validated test trials.

**Figure 3.**
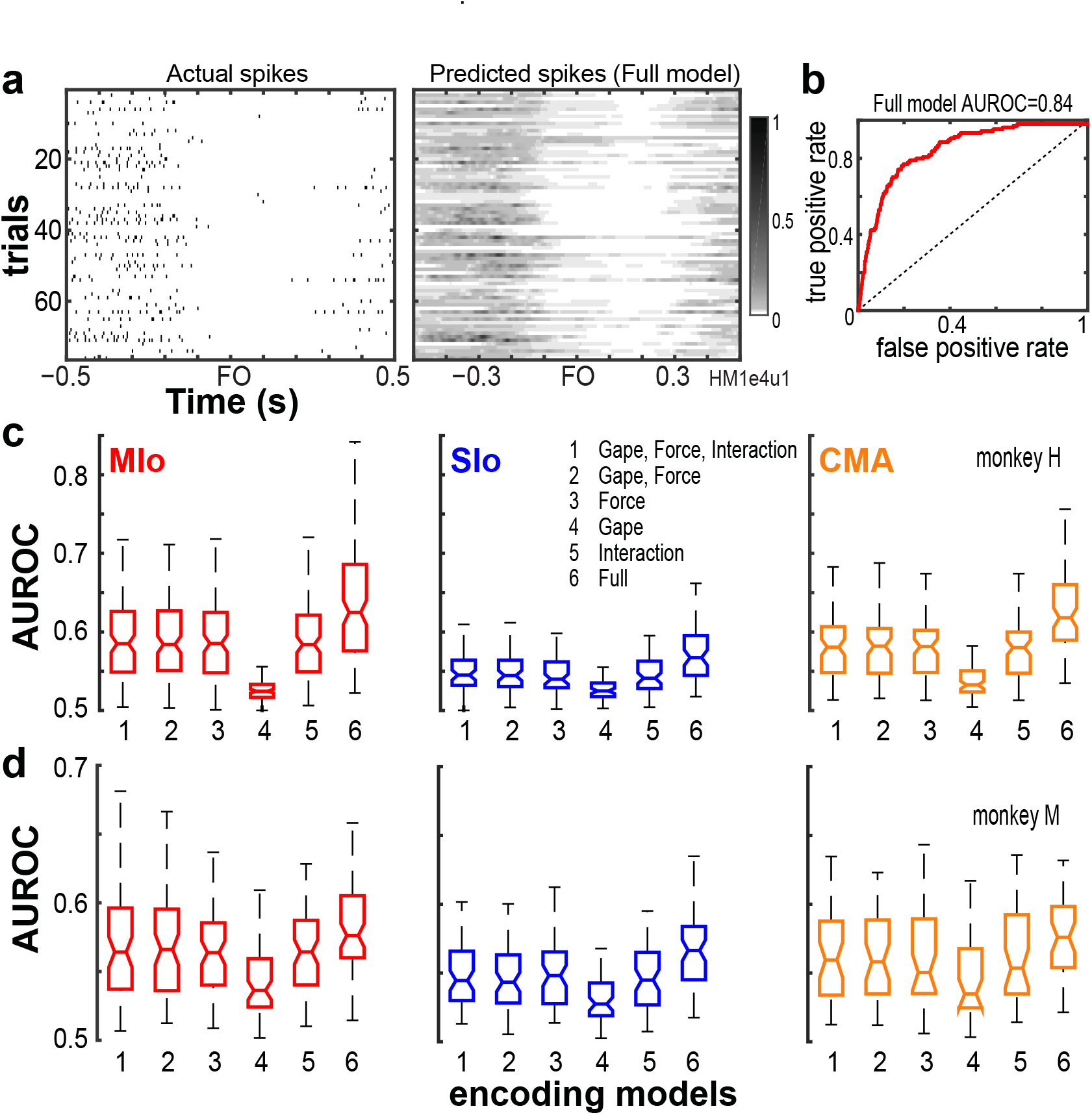
Performance of encoding models. (**a**), Actual firing rates of an example neuron vs. its predicted firing rates based on the full encoding model. (**b**), The full model’s goodness of fit was quantified using the AUROC on a cross-validated test data for the neuron shown in (a). Dashed line denotes chance level. (**c-d**), AUROC values from the population of neurons recorded from MIo, SIo and CMA are shown for each encoding model and animal. AUROC values are taken to be the mean across the 10-folds of cross-validation performed per neuron

We then compared the predictive power (using AUROC) of a full GLM model having all covariates with reduced models having only a subset of covariates. On average, we found that all input features used in all encoding models, full or reduced, were able to predict spiking activity of most neurons in MIo, SIo, and CMA significantly better than chance (Wilcoxon signed-rank test, monkey H: MIo: all *p*<4×10^−20^; SIo: all *p*<1×10^−11^; CMA: all *p*=8×10^−14^; monkey M: MIo: all *p*=2×10^−12^; SIo: all *p*=2×10^−11^; CMA: all *p*=5×10^−8^). However, the separate and combined ability of bite force, gape, and spike histories to predict the spiking of neurons differed; AUROCs were significantly different across the various encoding models, with the full encoding model showing the best performance and the gape-only model exhibiting the poorest performance (Fig. 3c-d, Kruskall-Wallis test, monkey H: MIo: all *p*=4×10^−44^; SIo: all *p*<5×10^−14^; CMA: all *p*<4×10^−25^; monkey M: MIo: all *p*=2×10^−8^; SIo: all *p*=6×10^−9^; CMA: all *p*=0.003).

We then sought to determine the relative importance of an input feature by comparing the performance of encoding models with and without the input feature in question. If an input feature contributes significantly to an encoding model, we would expect a model not to perform as well when that feature was removed. Figure 4 illustrates the degree of degradation of the predictive ability of an encoding model when either force or gape was removed by plotting each neuron’s AUROC against the encoding model that included both gape and force (intrinsic covariates were not included in the model for this analysis). When removing the force feature, a majority of neurons clustered above the unity line due to higher AUROCs in the combined force and gape model, indicating that excluding force from the encoding model degraded the model’s predictive ability (Fig. 4a-c, Wilcoxon signed-rank test, monkey H: MIo: *p*<5×10^−20^; SIo: *p*=2×10^−9^; CMA: *p*=6×10^−12^; monkey M: MIo: *p*=1×10^−8^; SIo: *p*=1×10^−6^; CMA: *p*=0.002). This was not the case when gape was removed as shown in most neurons clustering along the unity line (Fig. 4d-f, Wilcoxon signed-rank test, both monkeys, all areas, *p*>0.10). Thus, bite force was a more accurate predictor of spiking activity than gape. We also considered the possibility that the interaction between gape and force might contribute significantly to model performance beyond the combined contribution of gape and bite force. However, our results did not show any evidence for this; the model that included force, gape, and their interaction performed similarly to a model that excluded their interaction (Fig.3c-d, compare 1 vs. 2, Wilcoxon signed-rank test, both monkeys, all areas, *p*>0.10). Lastly, the full encoding model, that included force, gape, their interaction, and spike history, outperformed encoding models that did not include spike histories (Wilcoxon Paired sign rank test, monkey H: MIo: *p*=5×10^−22^; SIo: *p*=8×10^−12^; CMA: *p*=2×10^−14^; monkey M: MIo: *p*=3×10^−12^; SIo: *p*=8×10^−12^; CMA: *p*=1×10^−7^).

**Figure 4.**
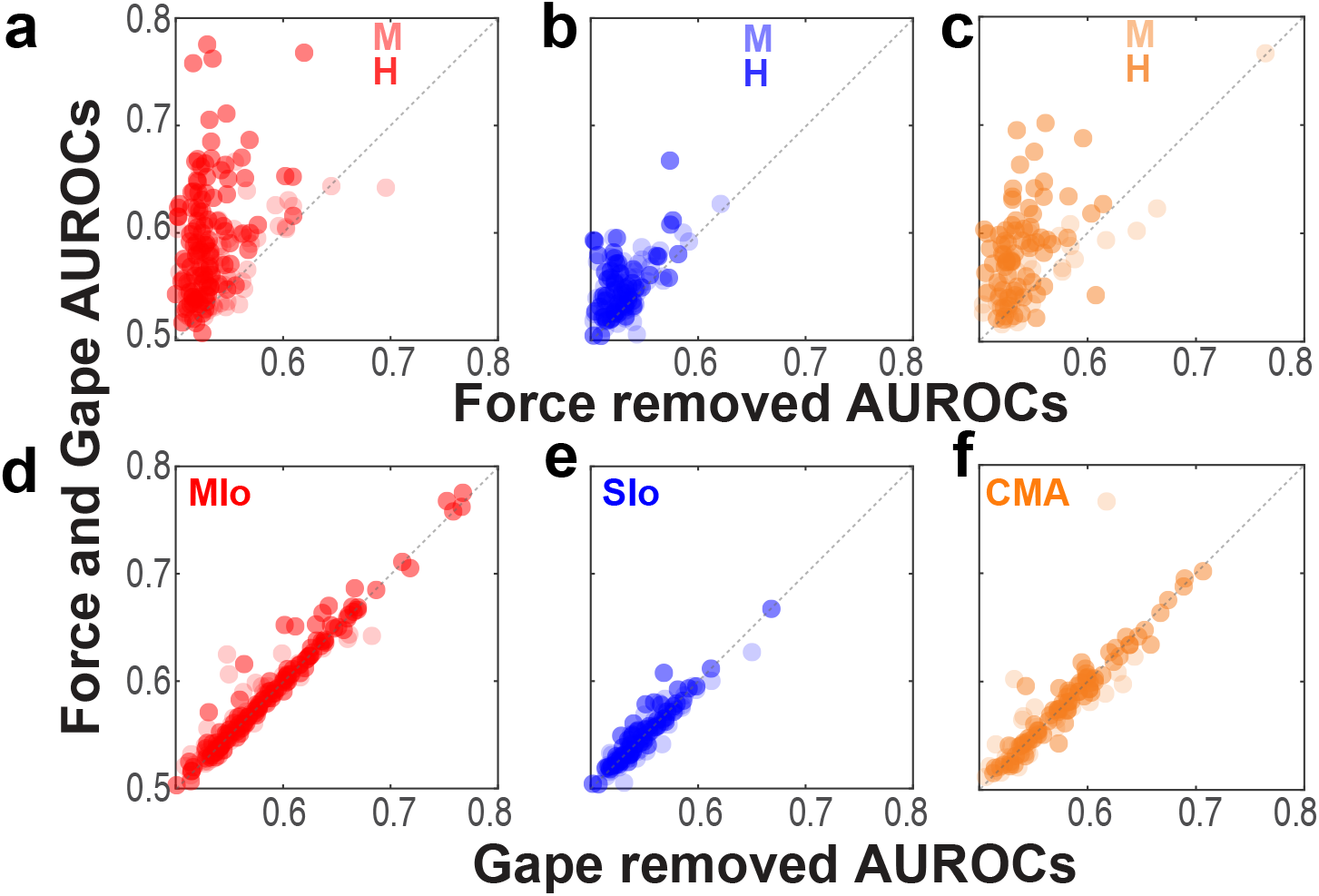
Comparison of model performance at an individual neuron level. (**a-c**), Relation between AUROCs of the joint force and gape model vs. the reduced model when force was removed. Each circle corresponds to a neuron’s AUROCs. Shown for each animal and for MIo, SIo, and CMA, respectively. AUROCs above the unity line (dashed line) denotes higher AUROCs in the joint vs. reduced model. (**d-f**), As in (a-c), but comparing AUROCs of the reduced model of when gape was removed.

While bite force accounted for most of the information that reduced encoding models used to predict spiking of individual neurons, the full encoding model that includes all input features (i.e., spike history, bite force, gape, and the interaction between them) outperformed all other reduced encoding models. Notwithstanding, each input feature carried distinct information capable of predicting spiking activity of neurons in OSMcx as shown by reduced encoding models performing above chance level.

### Distribution of neurons encoding bite force or gapes

We then evaluated whether there was any difference in the proportion of neurons encoding bite force compared with neurons encoding gape in the three studied areas of OSMcx. We identified ‘force-‘ or ‘gape-related’ neurons as neurons whose AUROCs in the force or gape only encoding model, respectively, were significantly higher than chance level (Wilcoxon signed-rank test, *p<0.05*). Force-related neurons, comprising a mean of 91% (SE 4%) of the recorded neuronal population across all areas and animals, were predominant over gape-related neurons (58%, SE 4%) (Fig. 5, *X^2^* test, monkey H: MIo: *p*=1×10^−16^; SIo: *p*=5×10^−6^; CMA: *p=*6×10^−11^; monkey M: MIo: *p*<5×10^−8^; SIo: *p*<2×10^−8^; CMA: *p*<0.09). There were no significant differences in the proportion of either force- or gape-related neurons between any two cortical areas (*X^2^* test, *p<0.017* after correction for multiple comparisons, monkey H: all *p*>0.10; monkey M: all *p*>0.04).

**Figure 5.**
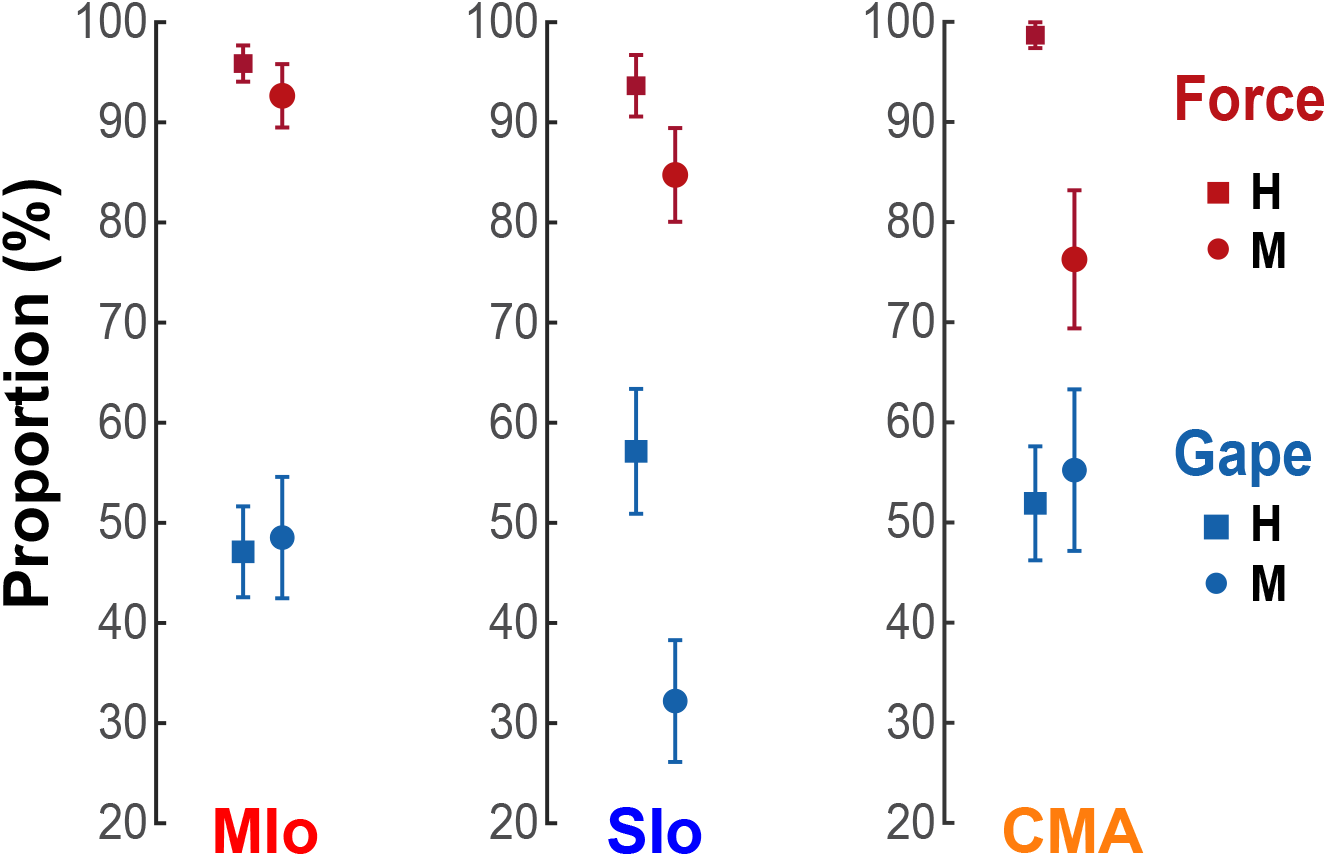
Distribution of neurons encoding bite force or gape. Proportion of neurons with AUROCs that were significantly higher than chance level (Wilcoxon signed-rank test, *p<0.05*) in the reduced force and gape models. Shown for each cortical area and subject. Error bars indicate±1 SEM (based on a binomial distribution assumption).

### Comparison of encoding model performance between cortical areas

To determine whether MIo, SIo, and CMA differ in the encoding of bite force and gape, we evaluated differences in the predictive ability of encoding models that included force only, or gape only, or both force and gape across the three areas. Using models with bite force and gape, we found a main effect of cortical area, where the activity of MIo was predicted better than the activity in SIo in both animals (Fig. 6a, Kruskall-Wallis test, monkey H: *p*=6×10^−9^; monkey M: *p*=0.0007, *post-hoc paired comparison with Bonferroni correction MIo vs. SIo, H: p*<0.0001, M: *p*<0.001) but not any better than activity in CMA (Fig. 6a, *post-hoc MIo vs. CMA*, *p>*0.10). When comparing encoding models that had bite force as the only predictor for spiking, the predictive ability of force in MIo was better than SIo but not any better than CMA (Fig. 6b, Kruskall-Wallis test, monkey H: *p*=5×10^−10^ *post-hoc MIo vs. SIo, CMA vs. SIo, p<*0.0001; monkey M: *p*=0.017, *post-hoc MIo vs. SIo, p<*0.05, *CMA vs. SIo, p*>0.10; both monkeys: *post-hoc MIo vs. CMA*, *p*>0.10). When encoding models had gape as the only predictor, a main effect of cortical area was observed in monkey H where the model’s predictive performance was best in CMA (Fig. 6c, Kruskall-Wallis test, monkey H: *p*=0.0016, *post-hoc CMA vs MIo, p*<0.01, *CMA vs SIo, p*<0.05). No significant differences between cortical areas were found in monkey M (Kruskall-Wallis test, *p*=0.055).

**Figure 6.**
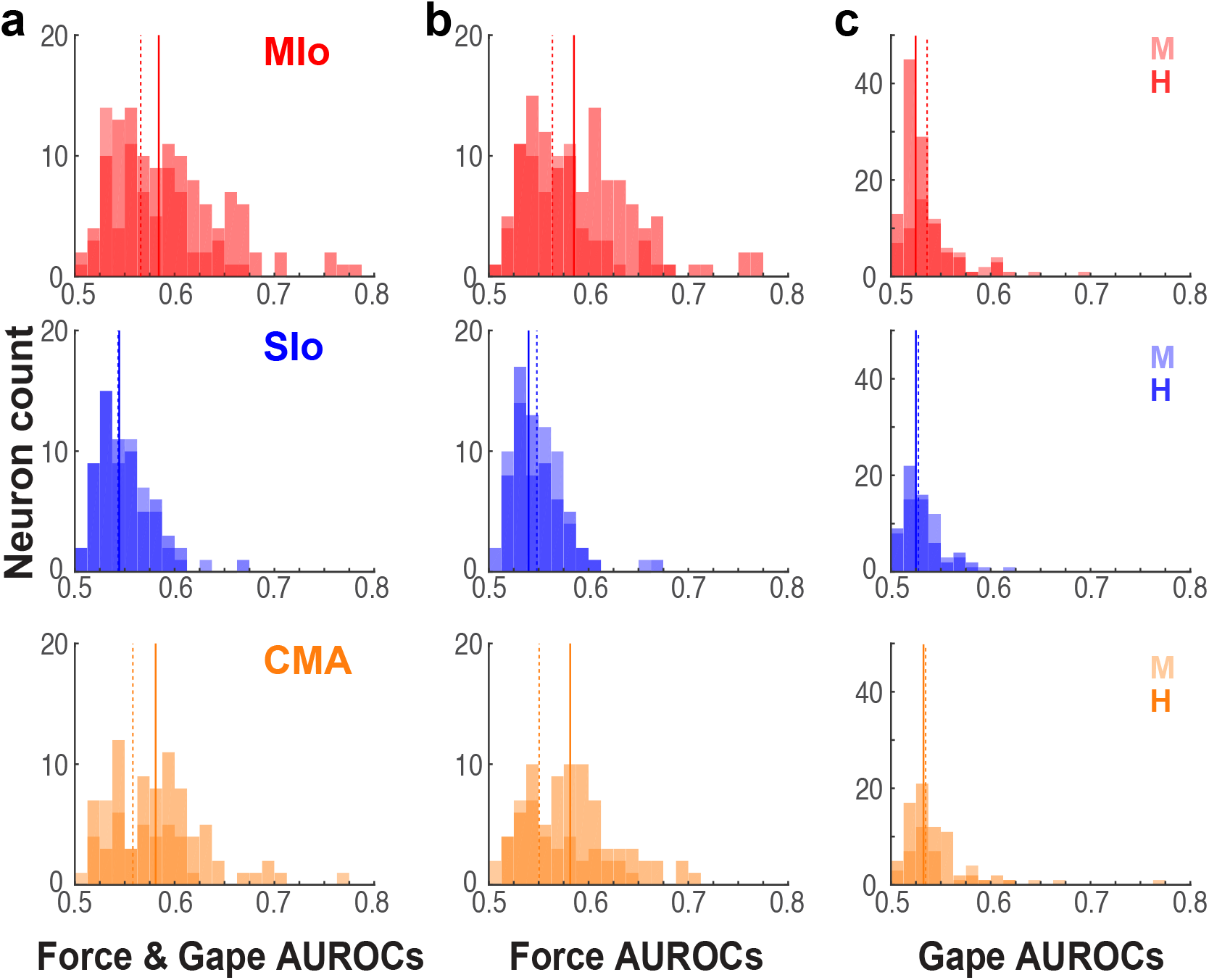
Performance of encoding models differed across cortical areas. (**a**), Distribution of significant AUROCs for the joint force and gape encoding model in MIo, SIo, and CMA, shown separately for each animal. Solid and dashed lines denote median AUROCs for monkey H and M, respectively. (**b-c**) As in (a) for AUROCs for the reduced force and gape models.

### Relative importance of temporal lags of bite force

Since we found that bite force is more strongly encoded than gape in all three cortical areas, we next evaluated the impact of temporal lags in bite force (from −156 ms to 208 ms relative to spiking) on predicting spiking of neurons in relation to bite force. For this analysis, we used the absolute values of the β coefficients of the temporal lags of bite force from the encoding model with force as the only predictor. Only β coefficients that were significantly different from zero were included (*t*-Test, *p*<0.05). For each neuron, we found the time lag that was associated with the largest β coefficient and computed the distribution of time lags across neurons for each cortical area (Fig. 7). Although the distributions of time lags were quite broad, there were important differences across cortical areas. The median time lags for MIo were 52 ms and 26 ms for monkeys H and M, respectively, indicating that force lagged spiking and consistent with the view that MIo drives force generation. In contrast, the median time lags for SIo were 0 ms for both monkeys. In CMA, the results were inconsistent across animals suggesting a more heterogeneous temporal relationship between force generation and neural responses (H:104 ms, M: 0 ms).

**Figure 7.**
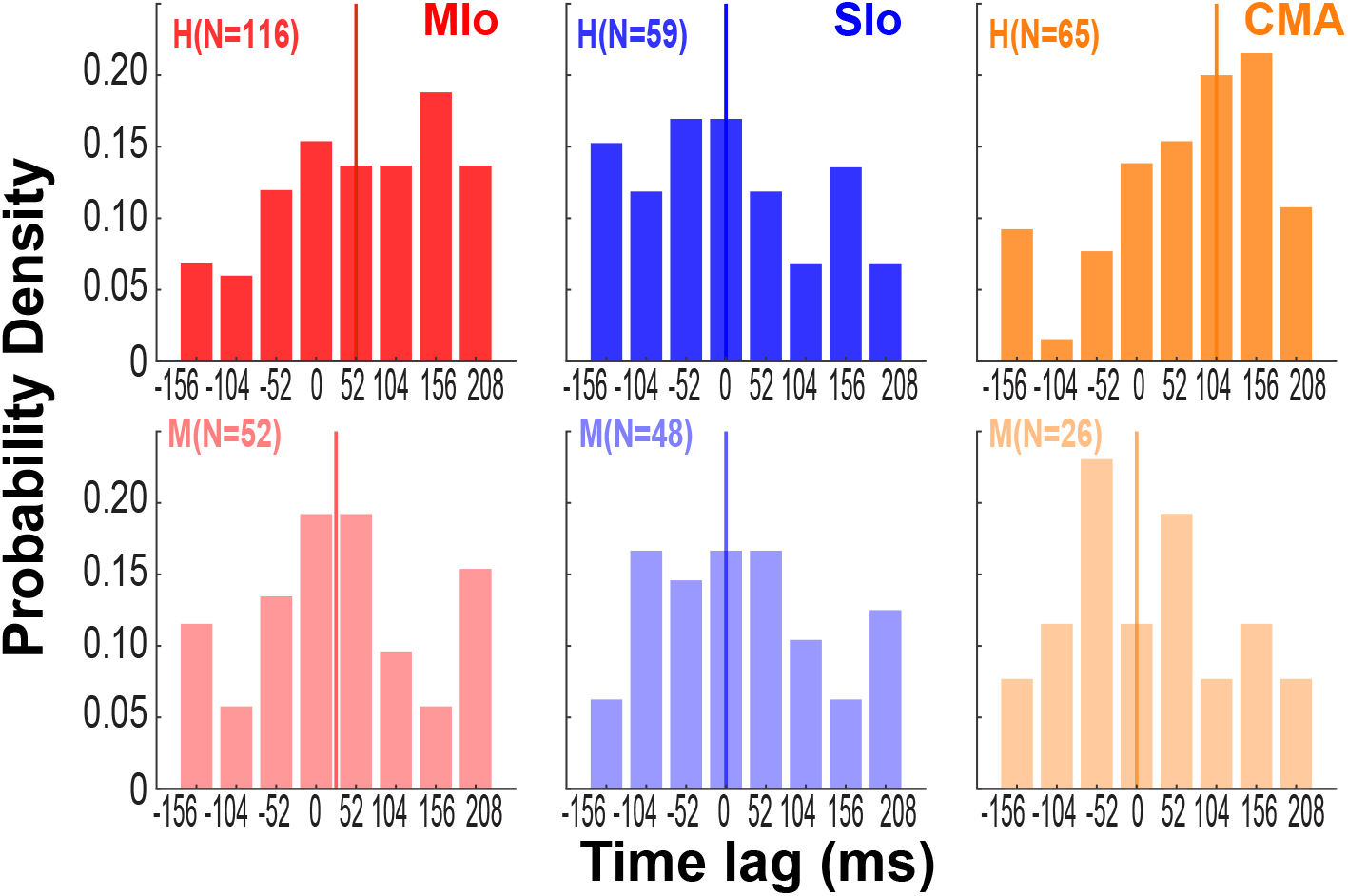
Preferred temporal lags of bite force differed across cortical areas. Distribution across neurons of time lags associated with maximum β coefficient for force in each cortical area and subject. Colored vertical lines indicate median values of each distribution.

### Relative importance of spike history timescale to spiking activity

The β coefficients for the most immediate spike history (16 ms) were higher than β coefficients for spikes that occurred further in the past (44 or 108 ms) for all areas in both animals (Fig. S3a-b, Kruskal Wallis, H: all *p*<2×10^−5^, M: all *p*<3×10^−5^, *Post-hoc* multiple comparison with Bonferroni correction, *p*<0.05). This indicates that immediate past history outweighs the other timescales in the ability to predict spiking of neurons in OSMcx.

### Population encoding of behavioral parameters

While encoding models using GLMs showed that bite force more accurately predicted the individual neuron’s spiking than did gape, our simultaneous, multi-site recordings allowed us to examine how activity at the population level in MIo, SIo, and CMA represents bite force and gape. Thus, we investigated how activity of neuronal populations in MIo, SIo, and CMA might distinguish between these behavioral parameters. Here, we used demixed principal components analysis (dPCA) (34) to decompose the dependencies of the population activity into a task-independent parameter of *time* (for activity related to the progression through the behavioral trial), and task-dependent parameters of *bite force* and *gape*, and the *interaction* between them. Figure 8 illustrates the cumulative variance in the population signal accounted for by demixed principal components (dPCs), i.e., neural modes, and the variance accounted for by individual task parameters for each cortical area. Over 70% of the variance was accounted for by 7-13 dPCs in monkey H (Fig. 8a) and 6-20 dPCs in monkey M. The first 5 dPCs showed very good demixing of task parameters as most of the component variance was explained by a single task parameter, such as the time-related activity for the first dPC or bite force for the second dPC in SIo (Fig. 8b center). While population activity could be decomposed into individual task parameters, the task parameters that explained most of the variance differed across cortical areas and as a function of the design of trial presentation (blocks of single gape in monkey H vs. gapes randomized trial-to-trial in monkey M, see Methods). First, time-related activity in all three areas was very prominent in monkey H. This accounted for most of the variance across cortical areas (45-60%). In contrast, time-related activity accounted for only a meager 8-27% in monkey M (pie charts in Fig. 8b-c, Fig. 8d). Second, variances accounted for by gape in all three areas were substantially higher when gapes were randomized trial to trial (20-57% in monkey M vs. 7-10% in monkey H, Fig. 8d). Moreover, explained variances of gape when trials were blocked were comparable across all cortical areas but differed substantially in the randomized design, with the explained variance in CMA being nearly triple that of MIo (MIo: 20%, SIo: 36%, CMA: 57%). Third, total variance explained by bite force was 2-3 times higher than variance explained by gape in monkey H (Fig. 8d). The opposite was found in monkey M; total variance explained by gape was 2-3 times higher than the variance explained by bite force in SIo and CMA, respectively (Fig. 8d). Lastly, the variance explained by interaction between gape and force in all cortical areas were substantial in both monkeys, suggesting a relative importance of the coordinated control of these two parameters (Fig. 8d). Bite force and the interaction were also modulated by task context, with substantial changes noted in MIo and SIo. Similar results were also found when dPCA was performed on a subset of trials and on two other datasets (Fig. S4).

**Figure 8.**
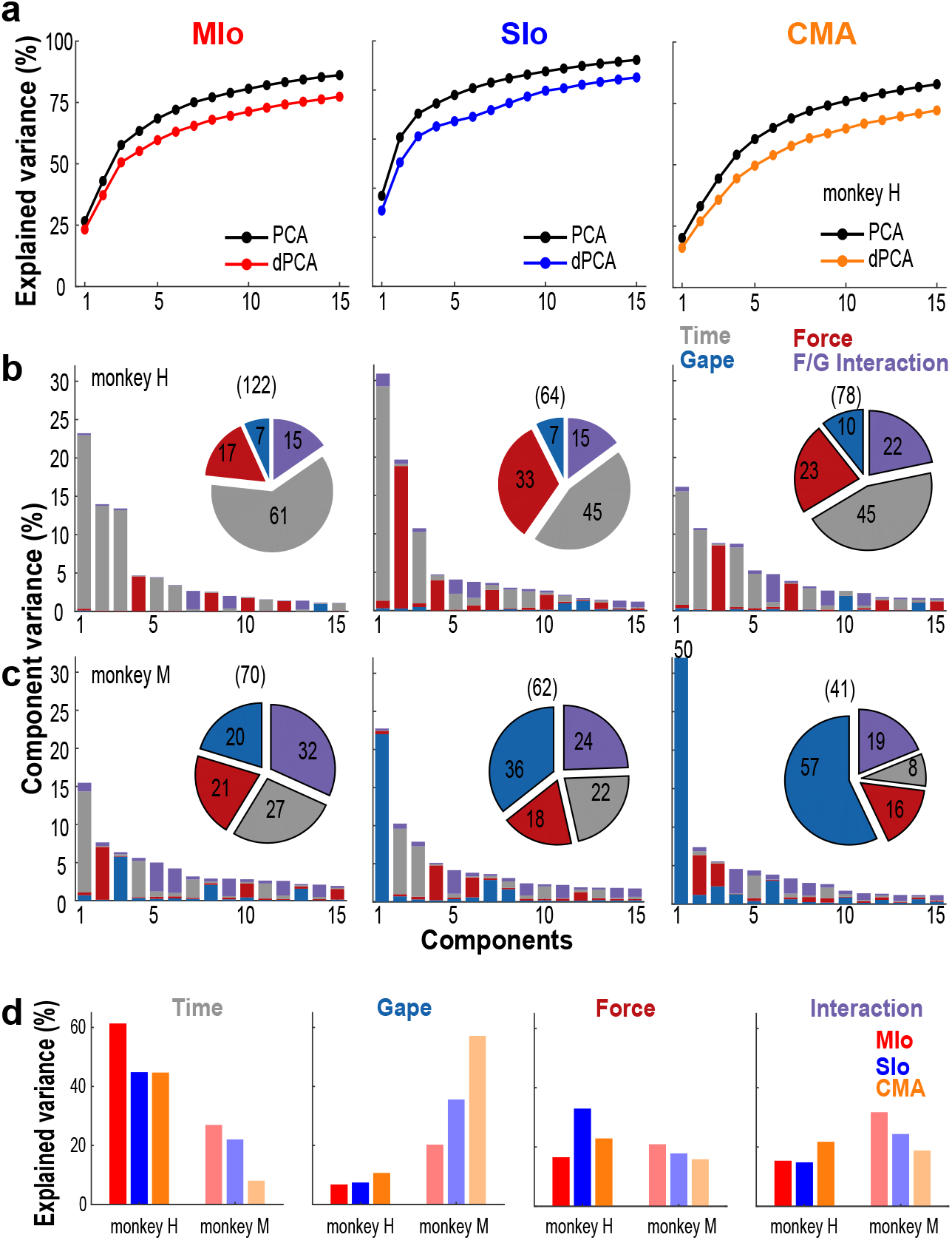
Neural variance accounted for by demixed principal components. (**a**), Comparison of cumulative variance explained by PCA and dPCA. Shown for the first 15 components and for each cortical area separately. Data from monkey H. (**b-c**) Bar graphs illustrating the proportion of variance accounted for by each task parameter (color) corresponding to individual dPCs in monkeys H and M, respectively. Single-colored bars depict complete demixing. Pie chart illustrates the proportion of variance (%) explained by task parameters. Numbers in parenthesis denote total number of neurons used in the analyses. Shown for each cortical area. (**d**), Across-area comparison of the proportion of variance (%) explained by each task parameter shown for both subjects.

Figure 9 illustrates the linear projections of population activity (i.e., latent activity) in MIo, SIo, and CMA corresponding to dPCs with the highest explained variance for each of the task parameters. Across cortical areas and animals, the task-independent, time-related activity, which captured the temporal progression of the population activity across the trial, was nearly identical across all trial types (i.e., combination of 3 levels of force and gape, Fig. 9a-b). The temporal evolution of the time-related activity exhibited maximal modulation around force onset.

**Figure 9.**
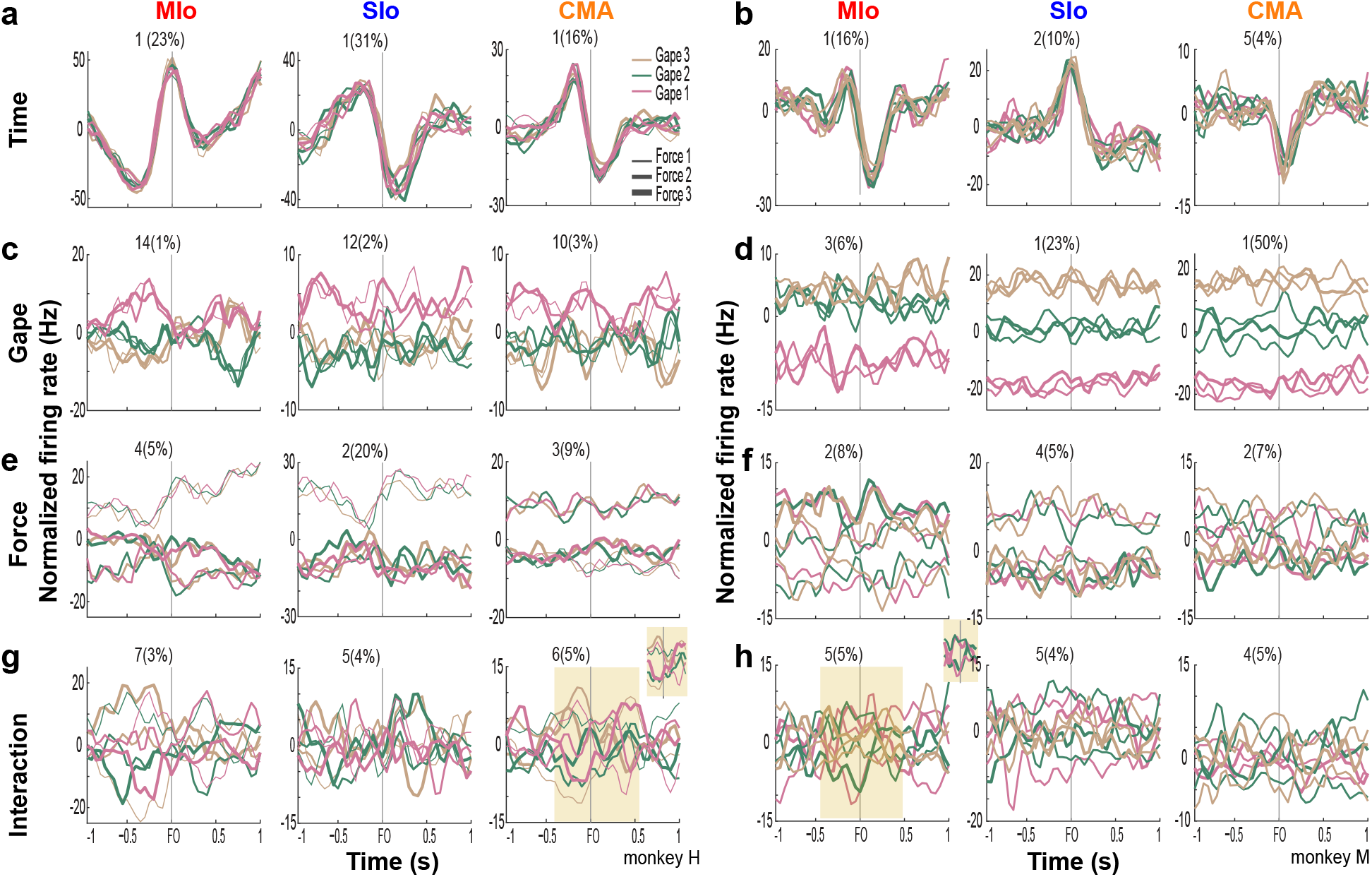
Latent activity of leading demixed principal components of task parameters. (**a-b**), Projections of population activity onto the leading dPCs of condition-independent parameter of time in monkeys H and M, respectively. Each subplot shows 9 lines corresponding to 9 trial types, the component number and corresponding explained variance. Shown for each cortical area. (**c-h**), As in a-b, for gape, bite force, and the interaction between gape and force.

The degree of separation of latent activity across gape distances differed between areas and subjects (Fig. 9c-d). In monkey H, gape-related latent activity in all cortical areas separated minimal gape from medium/wide gape (Fig. 9c). In monkey M, latent activity at all gapes was well-separated in SIo and CMA while activity of the lead dPC in MIo separated only between minimal gape and medium/wide gape (Fig. 9d). Across areas and animals, latent activities corresponding to different bite force levels were also separated, although tuning to bite force levels was observed at varying strength and times relative to force onset (Fig. 9e-f). Latent activity of all force levels was well-separated in MIo, with maximal separation occurring at force onset in both subjects. Both SIo and CMA exhibited distinct activities between two force levels only. Lastly, activity of dPCs corresponding to the interaction between gape and bite force showed varying degrees of separation at different times relative to force onset (Fig. 9g-h). For example, trial types were more separated around 0.3 s prior to force onset but became more overlapping after force onset (Fig. 9g inset). The activity of interaction components appeared complex, having distinct and overlapping activity patterns for certain gape-force combinations; low bite force generated at gape distances 1 and 2 had activity patterns opposite to high bite force applied at these gape distances (Fig. 9h inset). A subset of neurons that carry both gape and bite force information may underlie the coordination between these features.

The latent activity patterns of dPCs provided useful information about the modulation of population activity relative to behavioral events and task parameters, motivating us to evaluate the performance of dPCs in decoding gape and force at a single-trial level. Using the first dPC for each task parameter as a fixed linear classifier, we evaluated the accuracy of classifying gape and force levels. Classification of gapes was significant in monkey M only. Figure 10a shows significant classification of gapes ±1 s relative to force onset in SIo and CMA and in shorter periods in MIo. In the case of bite force, classification accuracy was significant in all areas for monkey H for most periods and in MIo and SIo for shorter periods in monkey M (Fig. 10b-c). Classification accuracy using interaction components did not show any time period with significant performance.

**Figure 10.**
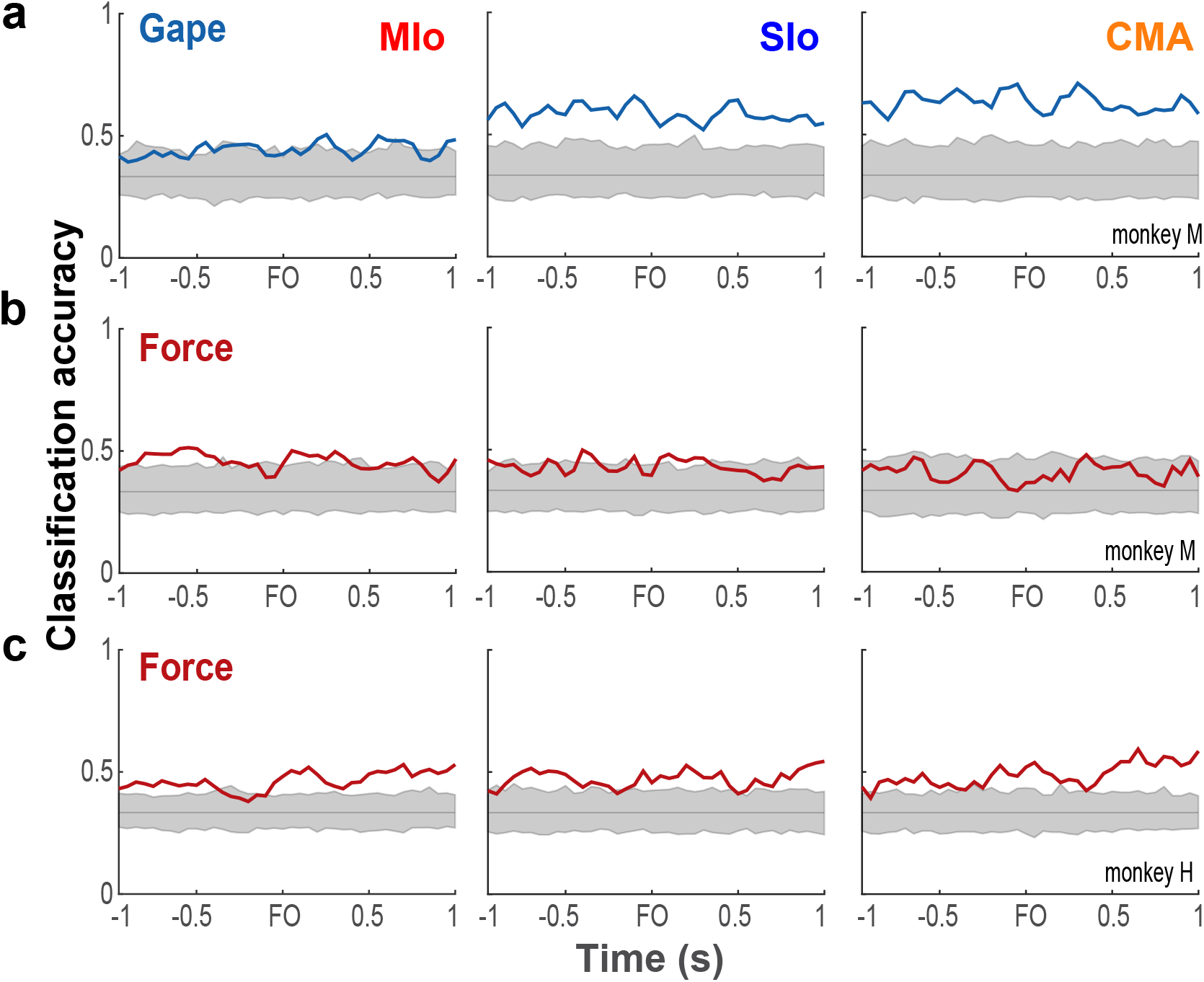
Classification performance of bite force and gape components. (**a**), Classification accuracies (blue line) of linear classifiers given by the first gape dPC in monkey M shown for each OSMCx area. Shaded gray area correspond to the distribution of classification accuracies expected by chance (solid gray line) as estimated by 500 iterations of shuffling procedure. (**b-c**), As in a but using the first bite force dPC in monkeys M and H, respectively.

## Discussion

In this study, we investigated how individual neurons and neuronal population in MIo, SIo, and CMA encode varying levels of bite force generated at varying gapes. To our knowledge, our study is the first to *(i)* investigate simultaneous encoding of bite force and gape, *(ii)* evaluated in three different areas of OSMcx, and *(iii)* at both the individual neuron and population levels. The activity of individual neurons in all three areas was strongly tuned to bite force albeit stronger in MIo than SIo. Population activity revealed robust tuning to gape when gape was randomized from trial-to-trial.

### Spiking activity of individual neurons in OSMcx is better predicted by bite force than by gape

Past studies demonstrated that MIo and SIo neurons modulate their activity to changes in bite force, jaw position, and movement (16, 35–37). In the current study, simultaneous recording in MIo, SIo, and CMA while subjects performed a biting task at varied combinations of gape and bite force levels allowed us to determine the relative importance of these task parameters in predicting the firing of neurons and to explore how these cortical areas might assume diverse roles in the control of a functionally important oromotor behavior. We demonstrated that when subjects performed a biting task, neurons in each of the three areas encoded bite force more strongly than gape (see Fig. 4) and that the force-related neurons were more predominant than the gape-related neurons. This was true regardless of the sequence used in presenting trials (i.e., blocked vs. randomized gapes). A possible explanation is that the task involved dynamic control of bite force whereas gape remained constant during a trial. The different control requirements for bite force and gape influence how muscles are activated, and which sensory information is more relevant. Because the task does not require voluntary control of jaw depression, activation of jaw depressors (anterior digastric, mylohyoid, and inferior head of the lateral pterygoid) is not expected because the lower jaw is passively depressed to a predetermined gape prior to the generation of the required bite force. Instead, jaw elevators (masseter, temporalis, medial pterygoid, and superior head of the lateral pterygoid) are expected to be activated to produce the required force level that varies during the trial. In this scenario, OSMcx neurons may be involved in the selective excitation of jaw elevators and inhibition of jaw depressors for the generation of varying bite forces (35, 38). The OSMcx receives information about the changing magnitudes of bite force and gape, in part via thalamus. This information is derived from neurons in the trigeminal mesencephalic nucleus and trigeminal ganglion that innervate muscle spindles of masticatory muscles or the mechanoreceptors of the periodontal ligaments of the teeth (11, 39–43). As sensory information on the position of the lower jaw remains unchanged during bite force generation, the most critical sensory information for successful task performance is the magnitude of applied bite force.

### Gape-related activity is better represented at the population level

Linear projections of population activity (i.e., latent activity) using dPCA in MIo, SIo, and CMA revealed robust tuning to gape that was not apparent at the level of individual neurons. Indeed, in monkey M where gape varied randomly trial-to-trial, the neural variance accounted for by gape was double to triple that of bite force (see Fig. 8d) and single-trial decoding of gape distances was significantly higher than chance (see Fig. 10a) notwithstanding the poor predictive ability (see Fig. 3d) of single neuron encoding models with only gape as the input feature. It is of interest that the temporal dynamics of the gape-related population activity revealed a cyclic or oscillatory pattern (Fig. 9c-d). These oscillations may be related to cyclic, short-range jaw depression-elevation, and thus, bite force generation, throughout the trial exhibited by monkey M. Alternatively, the cyclic pattern may be related to non-movement related factors such as posture maintenance involving coactivation or reciprocal inhibition of jaw-closing and jaw-opening muscles, similar to postural control processes during limb movements (44–47). In this context, the gape-related activity of OSMcx neurons may set the postural state of the jaw to the appropriate postural background for movement based on the expected sensory and motor consequences of the interaction of jaw dynamics and environmental factors on which bite force is generated and fine-tuned to meet task demands. Thus, sensory and motor systems are prepared for upcoming information from the external environment as well as from internal biomechanical changes. Similarly, randomization of gape on a trial-to-trial basis may have increased the demand for attention and reduced the predictability of task parameters, thus, diminished the ability to anticipate the appropriate sensorimotor response.

### Neuronal population encoding of task parameters reveals context-dependent modulations

Here we showed that the latent activity of populations of neurons in MIo, SIo, and CMA discriminated between task parameters, consistent with previous findings in other brain regions (34, 48). Our dPCA analysis also allowed us to capture features of population activity that were common or diverse. All three OSMCx areas shared neural modes (dPCs) with comparable task-independent, time-varying activation patterns while accounting for population variance in varying proportions. The differing population covariances across three OSMCx areas may reflect differences related to motor or sensory signals while the comparable temporal structure of activation patterns across areas may reflect shared network dynamics and/or inter-areal connectivity. We also found that population activity in all three areas varied with the context of task execution (i.e., blocked vs. randomized) wherein the topology of the motor behavior was preserved (i.e., generating bite force at varying gapes). When gape was randomized on a trial-to-trial basis, the explained variance for time decreased as the explained variance for gape more than doubled, and single-trial decoding of gape distances became significant. These results were not observed in monkey H who was presented with blocked trials of single gapes. While slight variations in the implantation sites of the multielectrode arrays may be a contributing factor (Fig. S1), we speculate that this effect is minor as the proportions of force- and gape-related neurons were comparable between subjects. Alternatively, the trial-to-trial randomization of gapes may have a stronger contribution because there is an increase in the demand to regulate changes in gape from trial-to-trial. This suggests that task context may adjust the contribution of relevant task parameters in determining the population activity, thus serving as a population encoding of differing contextual information for similar movement topologies. The results are consistent with previous findings showing context-dependent modulation of cortical encoding during texture discrimination in task vs. no-task conditions and grasping behavior with regular vs. irregular ladder wheels (49, 50).

### Diverse functions of orofacial cortical regions

Neuronal activity patterns, RF features, properties of evoked rhythmic jaw movements, and behavioral and cortical effects of ablations or reversible cold blocks of OSMcx differ across these three OSMcx areas (10, 13, 32, 33, 51, 52, 14, 19, 20, 27–31). Our results also demonstrate that MIo, SIo and CMA are all involved in the control of bite force and gape but differ quantitatively in their representation of these two parameters. While individual neurons in all three cortical areas encoded bite force more strongly than gape, MIo and CMA were better than SIo in predicting spiking activity based on bite force. The similarity between MIo and CMA may be related to anatomical overlap between borders of lateral MIo and CMA. The differences between cortical areas are unlikely to come from differences in RF properties as the RFs of neurons in all three areas are similar in having bilateral representations, though they are predominantly contralateral in SIo (7, 10, 11, 14). Thus, their difference may reflect differing functions with regards to motor- vs. sensory-related signals as well as density of network connections with other brain regions. For example, in addition to inputs from SIo, relevant sensory inputs also reach MIo and CMA via thalamo-cortical or cortico-cortical pathways. These findings point more to the role of SIo in modulating these types of behavior rather than generating them. The better representation of gape in CMA at the population level may be related to the involvement of CMA, which includes the lateral zone of MIo (11, 13, 14), in both rhythmic jaw movements as well as more elemental jaw-opening movements, all of which involve changes in gape. Moreover, the distinctive patterns of evoked rhythmic jaw movements described in previous studies suggest a role for distinctive masticatory patterns that can be attributed to input-output organization in these three OSMcx areas. Cortico-striatal and cortico-tegmental projections differ between MIo and CMA but have similar thalamo-cortical connections (14). Further studies are required to determine whether the diverse functions of these three areas could be more pronounced during the different stages of feeding behavior wherein distinctive masticatory, tongue, and swallowing patterns are naturally generated.

## Materials and Methods

### Subjects

Data were collected from two adult female rhesus macaques (*Macaca mulatta*), monkey H (7.5 kg) and monkey M (5.8 kg). All protocols were approved by the University of Chicago Animal Care and Use Committee and complied with the National Institutes of Health *Guide for the Care and Use of Laboratory Animals.*

### Behavioral task

Two naïve monkeys were trained to perform a behavioral task that approximates natural incisor biting behaviors requiring the generation of different levels of bite force at varying gapes (i.e., jaw depression distances) (Fig. 1a). The bite force plates were computer-controlled to open at one of three gapes prior to the start of a behavioral trial (Fig. 1b). The bite plates remained in that configuration for the entire length of the trial. Strain gauges bonded to the bite plates recorded the bite force produced by the teeth engaging the bite force plate. Nine combinations of required bite force (3 levels) and gape (3 distances) composed the nine different trial types. The presentation order of gapes was randomized in monkey M and blocked in monkey H. In blocked presentation, three force levels were randomly paired with a single gape before moving on to another gape. With training, both monkeys successfully generated the required bite forces at each of the three gapes (Fig. 1c). Detailed description of the task can be found in Supporting Information, Methods.

### Electrophysiology

Under general anesthesia, each monkey was chronically implanted with silicon-based arrays of 64 or 100 microelectrodes (BlackRock Microsystems, Salt Lake City, UT) in MIo, SIo and CMA of the left hemisphere (Fig. S1). The microelectrodes on the array were separated from their immediate neighbors by 400 μm and their length was 1.5 mm for arrays implanted in MIo and 1.0 mm for SIo and CMA. Implantation sites were verified based on surface landmarks and exhibited movements of the tongue or fingers evoked by monopolar surface stimulation of MIo (50 Hz, 200 µs pulse duration, 2-5 mA) during the surgical procedure. Signals from both arrays were amplified with a gain of 5000, simultaneously recorded digitally (16-bit) with a sampling rate of 30 kHz and hardware-filtered using a high-pass filter fixed at 1 Hz first, followed by a low-pass filter with 7.5 kHz cut-off (Grapevine, Ripple LLC, Salt Lake City, UT). Spike data streams were digitally filtered with a high-pass filter at 250 Hz. Spike waveforms were stored and sorted offline using Offline Sorter (Plexon, Dallas, TX). Data from array channels with no signal or with large amounts of 60 Hz line noise were excluded.

### Data analysis

Spiking activity of individual neurons recorded from MIo, SIo, and CMA was used in all neural analysis (Table 1). ***Generalized linear model (GLM).*** To determine the relative importance of bite force and gape in predicting the firing of neurons and to compare encoding properties among OSMCx areas, we used GLM to predict the time-varying spiking activity of each neuron. The GLM approach allows us to measure how well a model predicts the probability that a neuron fires a spike in a small sampling window (4 ms) based on different combinations of extrinsic covariates (i.e., bite force and gape) and intrinsic covariates (i.e., spike history). *Extrinsic covariates.* For gape, we used the gape distances which were adapted to the subject’s mandibular length (monkey H: 11, 14, 17 mm; M: 8, 11, 14 mm). For bite force, we included bite force magnitude at 8 different time lags from −156 ms (i.e., force leads spikes by 156 ms) to 208 ms (i.e., force lags spikes by 208 ms) in 52 ms steps relative to the spike sampling window. We used multiple time lags because multi-lag GLM models using kinematic features have been shown to provide higher predictive power than models that include only a single, optimal lag (53–55). We also included an interaction term for gape and bite force to evaluate whether encoding of the interaction between these two features was better than encoding of each feature separately. Thus, we used a total of 17 input features (1 gape, 8 forces, 8 interactions between gape and force) as extrinsic covariates. *Spike history.* The current spiking activity of a neuron might also be affected by its own spiking activity in the past due to intrinsic physiological properties such as absolute and relative refractory periods. Thus, we included the neuron’s spike history as an intrinsic covariate. To account for short (16 ms), medium (44 ms), and long (108 ms) time scale effects of the neuron’s own spike history, we filtered binary spike trains with raised cosine basis functions (Fig. S2). A log link function was used to relate the logarithm of the firing intensity (which is approximately equivalent to the spiking probability given the small 4 ms spike-sampling window) to a linear combination of covariates, expressed as:

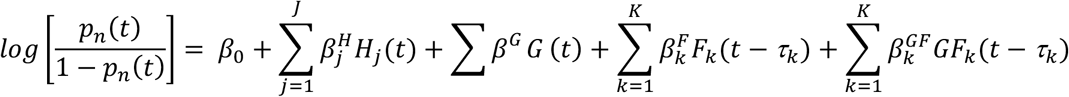

where *p_n_*(*t*) is the probability that neuron *n* fires a spike at time *t*, *β*_0_ represents the baseline probability that the neuron will spike, *H*_*j*_(*t*) is the value of the *j*^th^(of *J*) spike history timescale at time *t, G* is the gape distance at time *t*, *F_k_*(*t − τ_k_*) is the bite force at time (*t − τ_k_*), where *τ_k_* is the *k^th^* (of *K*) lead or lag time against the spike time at *t*, and *GF_k_*(*t − τ_k_*) is the interaction covariate at time *t − τ_k_*, and each covariate’s weight 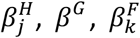 and 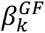, respectively. *Assessing the relative importance of each covariate.* We used different models based on the combination of input features used to predict a neuron’s firing. The full model includes all input features (bite force, gape, interaction, and the spike history of the neuron) while the reduced models have up to three of the input features removed (i.e. gape removed, bite force removed, and both force and gape removed, only force, only gape, only interaction, only spike history). To measure the goodness of fit of the encoding model, we compared the area under the receiver operating characteristic curve (AUROC) for 10 folds of cross-validated test data (i.e., 10 distinct sets of test trials that were not used to build the model) against chance level (53–57). ***Demixed Principal Components Analysis (dPCA).*** To investigate how activity of neuronal populations in MIo, SIo, and CMA might distinguish between behavioral parameters, we used dPCA (34) to decompose the dependencies of the population activity, ****X****, into components of time-dependent and task-dependent parameters: the task-independent parameter of *time*, *X_T_* (for activity related to the progression through the behavioral trial), the task-dependent parameters of *bite force*, *X_F_*, and *gape*, *X_G_*, and the *interaction* between them, *X_I_*:

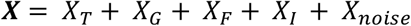

where ***X*** is the full data matrix with N rows of neurons which contain smoothed spike train of the *n*^th^ neuron for all task conditions and all trials. *X_T_*, *X_G_*, *X_F_*, *X_I_* are the linear decompositions of X into parameter-specific averages. dPCA then finds separate decoder (*D*) and encoder (*F*) matrices for each of these terms, *ϕ*, by minimizing the loss function: *L_dPCA_* = Σ*_ϕ_*‖*X_ϕ_* − *F_ϕ_D_ϕ_*X**‖^2^. To assess whether the condition tuning of individual dPCA components was statistically significant, we implemented the decoding method provided in the dPCA Toolbox (34) where classification accuracy was measured for each time point of a behavioral trial using the decoding axes of the first components of each marginalization, i.e., the first component of bite force was used to classify force levels, the first component of gape to classify gapes, and the first interaction component to classify all 9 trial types. The dPCA Toolbox uses cross-validation to measure time-dependent classification accuracy and a shuffling procedure to assess classification accuracy that is significantly above chance. We used 1000 iterations of stratified Monte Carlo leave-group-out cross-validation wherein on each iteration, one trial for each neuron in each condition was held out to form the test set and the remaining trials to form a training set. We used 500 iterations for the shuffling procedure.

**Table 1.** Number of neurons included in GLM and dPCA analyses.

|  | Monkey M | Monkey H |
| --- | --- | --- |
| Mlo | 70 | 123 |
| Slo | 62 | 64 |
| CMA | 41 | 78 |

We used the nonparametric Kruskal-Wallis one-way analysis of variance and the Bonferroni test for *Post-hoc* multiple paired comparison with significance level set at *P*<0.05, unless otherwise noted. All other analyses were performed using built-in and user-defined functions in Matlab (Mathworks, Inc.).

## Acknowledgments

This work was supported by NIH R01DE023816, NIH R01DE027236 and the University of Chicago Research Computing Center. We thank Dr. Kazutaka Takahashi for help with the Behavioral program set-up and the veterinary staff of the University of Chicago for animal care.

