## Supporting Information for "Multiple regions of primate orofacial sensorimotor cortex encode bite force and gape"

**This PDF file includes:**

Supplementary text
Figures S1 to S4

**Supporting Information Text**

**Supporting Methods**

***Behavioral task.*** The behavioral program was written using Matlab (Mathworks, Natick, MA) Toolbox Brain-computer interface to virtual reality (BCI2VR)(58). Behavioral event logs and timestamps were sent to the neural data acquisition system, Grapevine Neural Interface Processor (Ripple Neuro, Salt Lake City, UT). Force transducer (Micro Measurements, Raleigh, NC) analog signals were sampled at 1 KHz and stored using the neural data acquisition system. At the start of the recording session, monkeys were positioned such that their front teeth are pressed against the tip of the bite plates. The trial started with the movement of the bite plates to a specific gape. A cursor that represented the amplitude of the bite force applied on the transducer appeared 1 sec after the bite plate reached the desired gape. After a random period between 0.75 to 1.25 s from the appearance of the cursor, the base target window appeared to cue the monkey to keep the cursor within the base target window by applying a low-level isometric bite force for a hold period of 0.3 s. Upon successful hold at the base target, the force target window appeared to signal the monkey to move the cursor into the force target window. Three force target levels were set accordingly to the animal’s comfortable bite force range while keeping the visual displays of the target the same. The size of the force target windows corresponded to a range of ±2.5 units from the required force level. To achieve success, monkey had to generate the required force within the allotted time (5 s) and to hold the force 0.1 s for monkey M and 0.05 s for monkey H. To indicate success, the force target window changed color and the monkey received a reward. We set an inter-trial interval of 3 s.

**Supplementary Figures**

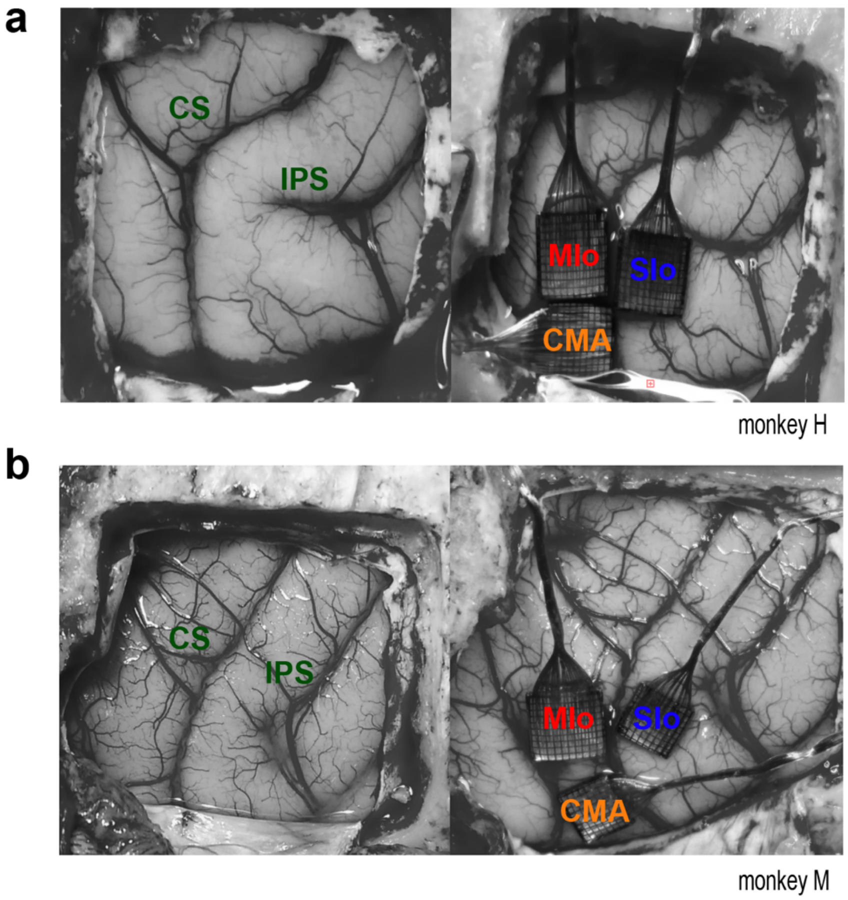


**Fig. S1. Anatomical location of cortical implants.** (**a-b**), Implantation sites of the microelectrode arrays in the left hemisphere of monkeys H and M, respectively (CS: Central sulcus, IPS: Intraparietal sulcus)

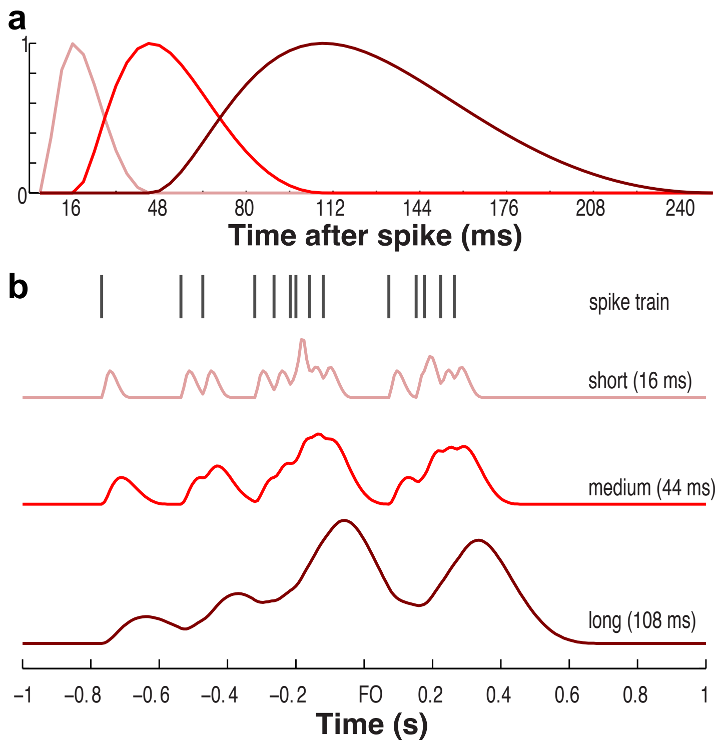


**Fig. S2. Spike history terms included in the encoding model.** (**a**), Three raised cosine basis functions of the form $b(t) = 0.5\cos(a\log(t+c) - \phi) + 0.5$, for $t$ such that $a\log(t+c) \in [\phi - \pi, \phi + \pi]$ and 0 elsewhere (59) were used for representing spike history filter. The peaks of the cosine curves on a log-transformed time axis represent short (16 ms), medium (44 ms), and long (108 ms) timescale effects of the neuron's own spiking activity. (**b**), Spike history terms. The original spike train of an example MIo neuron was convolved with each of the basis vectors shown in (a) to produce the three spike history terms used in the full encoding model.

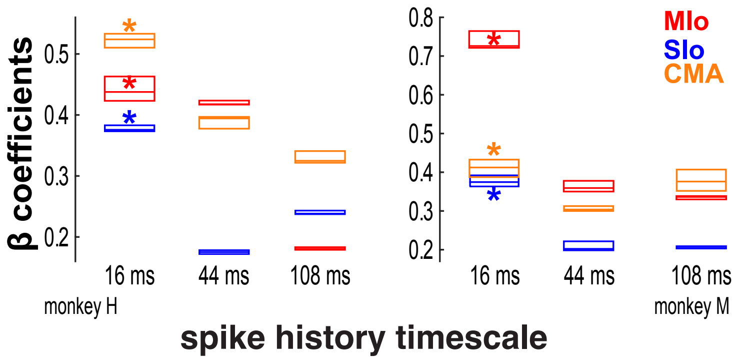


**Fig. S3. β coefficients of spike history.** Box plots of significant β coefficients of the spike history representing short (16 ms), medium (44 ms), and long (108 ms) timescales, shown for each area and animal separately. Asterisks denote spike history timescales that were significantly higher than other timescales within a cortical area ($p$<0.05).

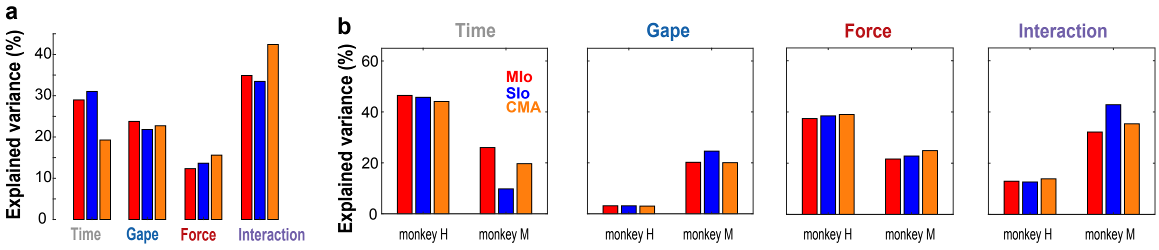


**Fig. S4. dPCA results using other dataset.** Percentages of explained variance when dPCA was performed on a subset of trials of the dataset of monkey M (**a**) and on other dataset of monkeys M and H.
